# *Fusarium graminearum* Ste2 and Ste3 Receptors Undergo Peroxidase Induced Heterodimerization

**DOI:** 10.1101/2023.09.14.557733

**Authors:** Tanya Sharma, Robert Y. Jomphe, Dongling Zhang, Ana C. Magalhaes, Michele C. Loewen

## Abstract

*F. graminearum Fg*Ste2 and *Fg*Ste3 are G-protein coupled receptors (GPCRs) recently shown to play roles in mediating fungal hyphal chemotropism and plant pathogenesis in response to activity arising from host-released peroxidases. Here, we follow up on the previous observation that chemotropism is dependent on both *Fg*Ste2 and *Fg*Ste3 being present at the same time; testing the possibility that this effect might be due to formation of an *Fg*Ste2-*Fg*Ste3 heterodimer. Initially the recombinant cell-surface expression of the *F. graminearum* GPCRs was validated in *S. cerevisiae* by confocal immunofluorescence microscopy. Bioluminescence resonance energy transfer analyses were subsequently conducted, where the addition of horse radish peroxidase (HRP) was found to increase the transfer of energy from the inducibly-expressed *Fg*Ste3-Nano luciferase (*Fg*Ste3-NLuc) donor, to the constitutively-expressed *Fg*Ste2-yellow fluorescent protein (*Fg*Ste2-YFP) acceptor, compared to controls. A partial response was also detected when an HRP-derived ligand-containing extract was enriched from *F. graminearum* spores and applied to the *S. cerevisiae* BRET system directly. The selectivity of the interaction was demonstrated by comparison to treatment with pheromones as well as an unrelated bovine GPCR, rhodopsin, fused to YFP as acceptor, that yielded no response when co-expressed with *Fg*Ste3-NLuc. Finally, the peroxidase-stimulated heterodimerization was validated by affinity pulldown. Taken together these findings demonstrate the formation of HRP and HRP-derived ligand stimulated heterodimers between *Fg*Ste2 and *Fg*Ste3. Outcomes are discussed from the context of the roles of ligands and reactive oxygen species in GPCR dimerization.

## 1 Introduction

G-protein-coupled receptors (GPCRs) represent the largest class of membrane proteins involved in cell signaling. They sense a broad range of environmental cues, referred to as ligands, and transduce these extracellular signals intracellularly. Classically, GPCRs form associations with heterotrimeric complexes consisting of hydrophilic guanine nucleotide binding (G-protein) alpha subunits (α), beta subunits (β) and gammas (γ) subunit (Calebiro & Godbole, 2018). Binding of ligands initiates conformational changes in the receptor leading to the dissociation of the α-subunit from the β- and γ-subunits, activating downstream signaling pathways.

Originally, it was thought that GPCRs functioned as independent entities. However, increasing evidence supports formation of functionally relevant receptor-receptor interactions. GPCR homodimerization and heterodimerization refer to the formation of complexes between two GPCRs of the same or different types, respectively. Both of these processes have been shown to be modulated by ligand binding, and in turn can modulate differential ligand binding, GPCR trafficking and signaling pathway preferences (Magalhaes et al., 2010, 2012; Marshall et al., 1999; Pello et al., 2008; Sleno & Hébert, 2018; Somvanshi & Kumar, 2012). These dimerization processes can occur on the membrane surface, or during protein synthesis and sorting in the endoplasmic reticulum. These receptor-receptor interaction events can change the cooperativity modulating ligand binding, where binding of one molecule of ligand influences the rate of binding of the other ligand molecule to the other GPCR, either positively or negatively (Sevlever et al., 2020).

Previous reports have highlighted the functional relevance of homodimerization and heterodimerization between GPCRs with homodimerization being a more commonly observed phenomenon (Angers et al., 2000; Breit et al., 2004). Approximately 27 GPCR heterodimeric receptor pairs have been characterized to date (Somvanshi & Kumar, 2012). In the case of the Class D α-factor pheromone GPCR, Ste2, it has been shown to homodimerize for transmission of α-factor pheromone dependent signalling (Shi et al., 2007), while evidence of heterodimerization is relatively sparse. One example of heterodimerization between GPCR subtypes includes the γ-aminobutyric acid (GABA) receptors, GABA_B_R1 and GABA_B_R2, where receptor functionality and cell sorting to the surface is dependent upon dimerization with each other (Marshall et al., 1999). Additionally, formation of heterodimers between entirely different receptors, for example bradykinin B2 and angiotensin AT1 receptors, can yield changes in the type of G-proteins they couple, from Gi as homodimers, to Gq as heterodimers (AbdAlla et al., 2000). This latter heterodimerization event has been shown to lead to alterations in endocytic properties in the cell.

Mechanistically, studies have shown dimerization-related structural rearrangements of transmembrane 4 (TM4) and TM5 as well as TM1 and TM7 to form the dimerization interfaces observed in GPCR dimer structures (Dijkman et al., 2018; Mondal et al., 2013). Additional interactions between the receptors can also be localized to either the N terminus (as observed for CCR5 homodimers) (Benkirane et al., 1997), C-terminal region (as seen in δ opioid homodimers) (Cvejic & Devi, 1997) or covalently linked dimers through cysteines located in extracellular domains (as reported for homodimerizing metabotropic glutamate receptor mGluR7 and the calcium sensing receptor CaR) (Bai et al., 1998; Romano et al., 1996). For Ste2 homodimerization a conserved GXXXG motif in TM1 was shown to be critical to dimer formation (Mueller et al., 2014; Overton et al., 2003; Shi et al., 2008). The GXXXG motif is a short, conserved sequence of amino acids commonly found in the transmembrane regions of many GPCRs, including TM3 of the Class D, a-factor pheromone GPCR, Ste3 (Overton et al., 2003).

Previously, we and others demonstrated the involvement of *F. graminearum Fg*STE2 and *Fg*STE3 GPCRS in fungal hyphal chemotropism toward the product of a catalytic reaction conducted by host-released peroxidases (Sharma et al., 2022a; Sridhar et al., 2020; Turrá et al., 2015). Δ*Fg*Ste2 and Δ*Fg*Ste3 deletion strains each individually lead to complete elimination of the chemotropic response to horse radish peroxidase (HRP). These individual deletions also both led to decreased pathogenicity on germinating wheat coleoptiles, underscoring the relevance of these receptors in mediating infection (Sharma et al., 2022a). Mechanistically, the receptor deletions individually led to decreased CWI-MAPK signaling compared to wildtype (WT) *F. graminearum*. Taken together, these findings highlight an enigma. How can two different receptors, each individually shown to modulate 100 % of chemotropic activity, show no activity in the absence of the other receptor? One possible explanation is that their activities are dependent on formation of a receptor heterodimer.

Here, we report the outcome of experiments testing the ability of *Fg*Ste3 and *Fg*Ste2 to form peroxidase-stimulated heterodimers. Using a recombinant expression system in *Saccharomyces cerevisiae*, a combination of biochemical and biophysical approaches including confocal microscopy, bioluminescence resonance energy transfer (BRET) and affinity pulldowns were applied demonstrating and validating the specificity of an observed *Fg*Ste2-*Fg*Ste3 interaction. The relevance of these findings to the molecular mechanisms underlying fungal chemotropism are discussed from the context of roles of receptor ligands and reactive oxygen species in GPCR dimerization.

## 2 Materials and Methods

### 2.1 Reagents D- (+)

Raffinose hydrate (Cayman chemicals,16773). All other reagents were purchased from Sigma™ unless stated otherwise.

### 2.2 Vector constructs and *S. cerevisiae* strain generation

Yeast centromeric vectors p415 (*LEU, GAL1* inducible promoter) and p416 *(URA3, GPD* constitutive promoter) were used to express chimeric proteins. The vectors were a kind gift from Barbara Montanini (University of Parma, Italy) (Corbel et al., 2017). The coding sequence for *F. graminearum* STE3 sequence was codon optimized for expression in *S. cerevisiae*, synthesized and cloned into the p415 vector using commercial service offered by Biobasics™. It was tagged at the C-terminus with an engineered deep shrimp luciferase, NLuc generating a fusion protein. This p415-*Fg*STE3-NLuc vector is referred to as ‘the Donor’. Similarly, *F. graminearum* STE2 coding sequence was subcloned into the p416 vector such that it was tagged with a C-terminal YFP, and thus p416-*Fg*STE2-YFP acted as ‘the acceptor’(Sridhar, 2023). Briefly, for generating fusion protein DNA constructs, the cDNA sequences were amplified without stop codons along with the respective reporter tags, NLuc for *Fg*Ste3 and YFP for *Fg*Ste2. They were followed by a glycine and serine rich linker molecule sequence (protein sequence: GGGGSGGGGS) that allowed spatial flexibility. This was followed by a FLAG tag (amino acid sequence DYKDDDDK) and stop codon.

These constructs along with a number of others, were either expressed alone or were co-expressed in select combinations to create the Test, Positive control and Negative control *S. cerevisiae* strains needed for BRET analysis (**Table 1)**. Three notable Negative controls were employed. First the *Fg*Ste3-NLuc donor was expressed alone to account for any bleed through signal arising from the donor into the acceptor wavelength emission (Donor alone). Secondly, *Fg*Ste3-NLuc was co-expressed with an un-related GPCR, bovine rhodopsin, fused to YFP referred to as Rho-YFP (Loewen et al., 2001) to produce the Negative (Rho) strain. Third, *Fg*Ste3 and *Fg*Ste2 were co-expressed in the absence of either of the two reporter fusions to account for background fluorescence, referred to as the Negative (NR (no reporters)) strain.

**Table 1:** List of yeast plasmids used in the study and *S. cerevisiae* strains generated upon co-transformation.

|  |  |
| --- | --- |
| **Vectors** | **p415 GAL1: *LEU2*, GAL1 inducible promoter** |
| 1 | *FgSte3*-L-NLuc-L-YFP-FLAG-Stop |
| 2 | *FgSte3*-L-NLuc-FLAG-Stop |
| 3 | *FgSte3*-FLAG-Stop |
| **Vectors** | **p416 GPD: *URA3*, GPD constitutive promoter** |
| 4 | *FgSte2*-L-YFP-L-NLuc-FLAG-Stop |
| 5 | *FgSte2*-L-YFP-FLAG-Stop |
| 6 | *FgSte2*-FLAG-Stop |
| 7 | Rho-L-YFP-FLAG-Stop |
| **Co-transformed vectors** | **STRAIN NAMES** |
| 1&4 | Positive |
| 2&5 | Test |
| 3&6 | Negative (NR; no reporters) |
| 2&6 | Donor alone |
| 2&7 | Negative (Rho) |

Fungal strain BY4741 YFL026W (*MATa his3*Δ*1 leu2*Δ*0 met15*Δ*0 ura3*Δ*0*) was grown in 50 mL of Yeast extract-Peptone-Dextrose (YPD) media and was used for transformation using the lithium acetate method as described with minor modifications (Gietz & Schiestl, 2007; Shi et al., 2009). Salmon sperm DNA (2mg/ml) was used as the carrier plasmid. The BY4741 YFL026W cells were transformed with the appropriate plasmids and the colonies were allowed to grow on SR/-Ura/-Leu plates (Synthetic Raffinose medium) for 2-3 days. For growth of transformants, single colonies were picked and individually grown in liquid SR/-Ura/-Leu at 29 *°*C.

### 2.3 Confocal microscopy for studying Ste3-NLuc and Ste2 YFP expression and localization

The yeast ‘Test’ strain co-transformed to contain vectors encoding the NLuc and YFP tagged receptors was grown in SR/-Leu/-Ura media overnight and then induced with 2 % galactose. The obtained cell cultures were treated with 4 μM HRP enzyme for 10 minutes (min) followed by centrifugation and subsequent washing in ice cold 1X Phosphatase Buffer Saline (PBS) consisting of 137 mM NaCl, 2.7 mM KCl, 10 mM Na_2_HPO_4_, 1.8 mM KH_2_PO_4_. They were fixed using 4 % paraformaldehyde for 20 min at 4°C. To enable antibody-based visualization of NLuc, Zymolase was added to the cell suspension for 1 hour (h) to digest the yeast cell wall. Following washing (3 times in PBS), cells were permeabilized with PBS containing 0.01 % Triton X-100 for 20 min. To reduce non-specific binding, cells were incubated in a blocking solution with PBS containing 1mg/mL Bovine Serum Albumin (BSA) for 30 min. Subsequently, cells were incubated with a 1:100 dilution of primary mouse anti-NLuc antibody (Promega, cat#7000) in PBS overnight at 4 °C. Following washing (3 times in PBS), cells were incubated with a 1:500 dilution of secondary rabbit IgG-antibody conjugated to Alexa fluor 647 (Thermo Fisher, A-21235) for 60 min at room temperature in the dark. Cells were added on top of the slide with a small amount of buffer, an anti-fading Mowiol media was added around the cells drop and a coverslip added on top.

### 2.4 Bioluminescence resonance energy transfer assay

This is a proximity assay based on radiative transfer of energy between Nano luciferase (NLuc) and yellow fluorescent protein (YFP), fused to *Fg*Ste3 and *Fg*Ste2 receptors respectively. NLuc was employed as the donor and YFP the acceptor construct as previously documented here (Corbel et al., 2017). Employing NLuc had the advantage of its smaller size and good separation between the NLuc and YFP emission spectra. Upon addition and oxidation of Nano-glo luciferase substrate (Promega, N1110) by NLuc, light is emitted with a peak at 480nm. YFP absorbs this if it is within the permissive distance (< 100 Å) of the NLuc light emission and in turn emits fluorescence at with a peak at 530 nm.

Transformed fungal strains were grown for 2 days at 29 °C in SR/-Ura/-Leu media to O.D 0.7-1 (density of 2 × 10^3^ cells per mL). Equal volume of culture (50 μL) was loaded in each well of a 96-well PerkinElmer ™ white optiplate, to which galactose was added at a final optimized concentration of 2 % (see below). The cells were incubated for 3 h at 29 °C for induction of the donor *Fg*Ste3-NLuc protein. Prior to recording the BRET reading, the cells were treated with HRP (final concentration 4 μM) dissolved in PBS and incubated at room temperature (RT) for 2 min. In cases where pheromones were applied, they were applied at a final concentration of 1 μM. Following this, 20μL of freshly diluted Nano-glo Luciferase Assay substrate (Promega ™, Madison USA, N1110) diluted to a ratio of 1: 1000 times in PBS was added. The relative luminescence units (RLU) readings were then acquired using a Spectra max M5^e^ multi-mode plate reader using the following emission filter settings for recording luminescence: NLuc: counting time 5 seconds; emission filter 480nm (±10nm) and YFP: counting time, 5 seconds; emission filter 530nm (±10nm). Measurements were performed on each well over time to measure the progress of reactions. Background luminescence detected in i) wells containing donor alone transfected cells (expressing *Fg*Ste3 fused to NLuc and *Fg*Ste2 alone) and ii) the Negative (NR) strain (expressing *Fg*Ste2 and *Fg*Ste3 alone and lacking NLuc and YFP fusions was subtracted. The experiments were performed with a minimum of three and maximum of ten replicates for each condition and repeated at least twice. The average of two independent experiments are represented in each graph. The BRET ratio was calculated by dividing the signal measured at 530 nm (YFP as acceptor) by the signal measured at 480 nm (NLuc as donor) after background emissions were subtracted as detected in Negative (NR) and Donor-alone strains. The netBRET was calculated by subtracting the ratio obtained for donor alone strain from the other strains being tested.

Expression levels of YFP tagged acceptor proteins were assessed using λex = 513 nm, λem = 530 nm prior to performing luminescence readings to make sure they were consistent across the samples and replicates. For quantification of the levels of donor protein, we used the sum of bioluminescence emission signal recorded from donor NLuc and acceptor YFP proteins as described previously (Besson et al., 2022). Z factor values were calculated (>0.5) to optimize the feasibility of the assay for subsequent conditions tested (Zhang et al., 1999).

### 2.5 Preparation of chemotropic ligands and applications to the BRET assay

The HRP-derived ‘ligand’ was obtained by treating 10 million *F. graminearum* spores with 0.05 μM concentration of HRP for 6 h at 28 °C. The spores were centrifuged at 6000 x g and the aqueous phase was collected and HRP was heat deactivated by incubating it at 95 °C for 10 min as demonstrated previously (Sharma et al., 2022b), leaving only the spore-derived product of the HRP reaction referred to herein as the ‘ligand’, and not HRP itself. The chemotropic activity of the ligand was confirmed in chemotropism plate assays (Sridhar, 2023), before being applied to the BRET assays. In other experiments, intact HRP or pheromone itself was applied directly to the BRET assays. The correlation between concentration of ligand (or HRP, or pheromone) and receptor BRET saturation was studied by assaying increasing concentrations of the stimulants. HRP was assayed from 0-10 μM, pheromones (a-factor and α-factor) were assayed at 1-10 μM, while the ligand, which could not be quantified, was applied at increasing volumes from 0-10 μL.

### 2.6 Optimization of galactose concentration for expression from p415-*Fg*STE3-NLuc donor vector

A saturation assay was carried out for determining the galactose concentration at which donor saturation was observed (BRET_max_). Samples (50 μL) of the yeast ‘Test’ strain culture were loaded in each well of a 96-well plate to which increasing concentrations of galactose were added from 0.01 to 8 % (in triplicate). The plate was incubated for 3 h (after optimization of induction time (see below)) at 29 °C for induction of *GAL1* promoter driving *Fg*Ste3-NLuc (donor) protein expression. HRP (4 μM) was added followed by 20 μL of Nano-glo luciferase substrate (Promega, N1110) to each well. The luminescence data was collected as described above. The BRET_50_ value was calculated and used going forward as the optimized concentration of galactose to be used for performing BRET assays. The expression of *Fg*Ste3-NLuc donor protein produced was also validated performing immunoblotting using anti-Nluc antibody (Promega cat#7000), see details below.

### 2.7 Temporal dynamics of heterodimer formation

With inducible expression of *Fg*Ste3-Nluc donor protein and constitutive expression of *Fg*Ste2-YFP acceptor protein ensuring that the receptors were expressed on the surface, the observed BRET responses were relatively fast. The BRET response was measured using a kinetic assay over a period of time ranging from 0 min to 1 h after HRP or pheromone or ‘ligand’ and luciferase addition. Yeast cells expressing the Test constructs were grown for 2 days in SR/-Leu/-Ura media and induced to 2 % galactose concentration for 3 h. The cells (50 μL) were pipetted in each well of a white optiplate (Perkin Elmer) and assayed with 4 μM of HRP and 20 μL of Nano Glo luciferase substrate (Promega, N1110) diluted to 1:1000.

### 2.8 *Fg*Ste2 and *Fg*Ste3 interaction by tandem-affinity pulldown

The yeast Test strain, as described for BRET in **Table 1** was used. The cells were grown for two days in SR/-Ura/-Leu media in 2 % galactose and exposed to HRP for 10 min. The peroxidase treated cells were pelleted by centrifuging at 4000 x g for 10 min, resuspended in 5 mL of PBS and lysed by vortexing with glass beads following a previous protocol with minor modifications(Hauser et al., 2007). All the steps were performed at 4 °C unless indicated otherwise. The solubilized membrane proteins were diluted 1:1 in dilution buffer (10 mM Tris/Cl pH 7.5, 150 mM NaCl, 0.5 mM EDTA) and incubated with GFP trap beads (ChromoTek; AB_2631357; with 95 % efficiency to bind YFP tagged proteins) with end-over-end mixing overnight at 4 °C. The beads were washed three times in wash buffer (10 mM Tris/Cl pH 7.5, 150 mM NaCl, 0.05 % Triton-X, 0.5 mM EDTA) and allowed to settle by gravity each time. The supernatant was removed. Bound proteins were extracted from the beads using 50 μL of acidic elution buffer (200 mM glycine pH 2.5). SDS sample buffer (2X concentration, 10 μL; 120 mM Tris/Cl pH 6.8, 20 % glycerol, 4 % SDS, 0.04 % bromophenol blue, 10 % β-mercapta-ethanol) was added and samples were incubated at RT for 15 min. Solubilized proteins were resolved by SDS-PAGE (10% acrylamide) and transferred to nitrocellulose membrane using standard wet electroblotting protocol. The blot was probed with 1:1000 dilution of anti-NLuc antibody (Promega cat#7000) and the bands detected by chemiluminescent detection.

### 2.9 Statistical Analysis

All BRET experiments consisted of 3 minimum technical replicates and were replicated at least two times. One-way ANOVA and multiple mean separation modules of the Graph pad prism (version 9.0, Graphpad Software, San Diego, CA) were used to calculate statistical differences and compare the means, respectively. Curve fitting was performed using a non-linear regression equation assuming a single binding site. Multiple comparisons of the data were performed by ANOVA.

## 3 Results

### 3.1 Recombinant expression of *Fg*Ste3-NLuc and *Fg*Ste2-YFP in *S. cerevisiae*

To confirm the recombinant expression of the *Fg*Ste3-NLuc and *Fg*Ste2-YFP fusion constructs, confocal imaging of the ‘Test’ strain (**Table 1**) was performed. For *Fg*Ste2-YFP, whose expression was under the control of a constitutive p416 GPD promoter, a robust signal was observed in green (**Figure 1A & 1B**). For detecting *Fg*Ste3-NLuc which was under the inducible p415 *GAL1* promoter, indirect immunofluorescence was employed. Following yeast cell wall digestion and permeabilization, anti-NLuc antibody followed by IgG secondary antibody conjugated to the fluorophore Alexa Fluor 647 (red color) were sequentially added to the cells. Significant red was observed concentrated at the shmoo regions (Figure 1A) and a more uniform, faint distribution was observed within the cell (Figure 1B). This distribution is consistent with previously reported localization of *Sc*Ste3 in *S. cerevisiae* (Davis et al., 1993). The imaging experiments were performed in conditions where receptors are not overexpressed (validated by BRET – see next section). Negative control (NR) strain (**Table 1**) treated as described above, accounted for the autofluorescence background signal (**Figure 1C**). The heterogeneity of the cell population with cells expressing only the *Fg*Ste2-YFP, only the *Fg*Ste3-NLuc and some expressing both receptors are highlighted (**Figure 1D**).

**Figure 1.**
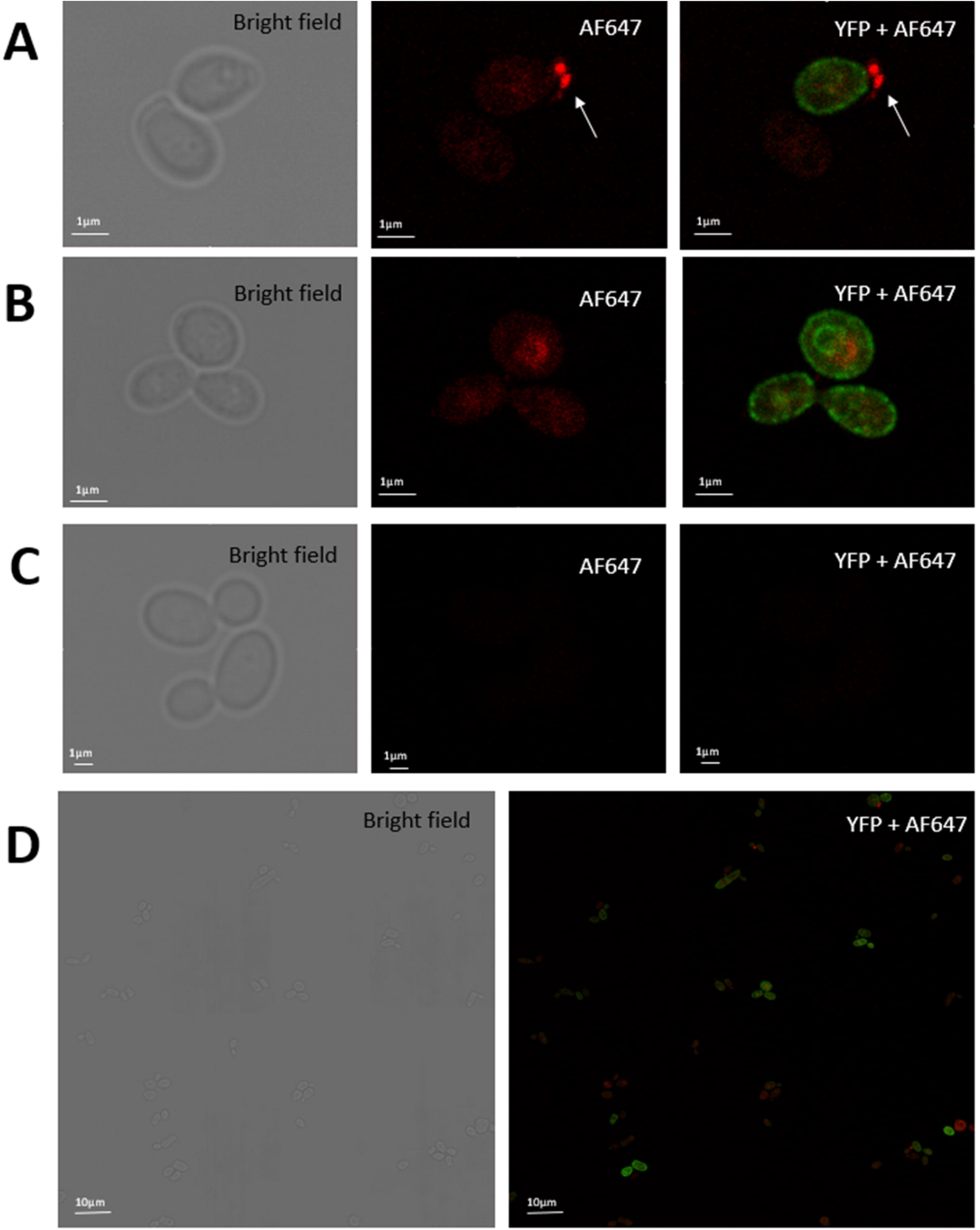
Expression of *Fg*Ste3-NLuc and *FgS*te2-YFP in *S. cerevisiae*. Confocal microscopy analysis (3.1x zoom) of the BRET Test strain co-expressing *Fg*Ste3-Nluc and *Fg*Ste2-YFP. Expression of the donor *Fg*Ste3-NLuc was detected using immunofluorescence with AF647 and is shown in red and the *Fg*Ste2-YFP is shown in green. **A)** *Fg*Ste3-Nluc labeled with AF647 concentrated around the shmoo region depicted with a pointed arrow. **B)** *Fg*Ste3-NLuc and *Fg*Ste2-YFP co-expressed in S. cerevisiae cells. More specifically, the receptor *Fg*Ste2-YFP localized mainly on the cell membrane, whereas the donor *Fg*Ste3-Nluc resided in cytosol. **C)** Negative control showed no fluorescence signals in both YFP and AF647 channels. **D)** Heterogeneous population of transformed cells in 2x2 Mosaic showing co-expression of *Fg*Ste3-NLuc and *Fg*Ste2-YFP in cell population. NLuc refers to Nano luciferase and YFP refers to Yellow Fluorescent protein.

### 3.2 Optimization of BRET assay parameters

Initially galactose-induced *Fg*Ste3-NLuc donor expression levels in *S. cerevisiae* were optimized. Galactose saturation assays showed a hyperbolic curve (**Figure 2A**). With an inducible *GAL1* promoter employed to drive the expression of the donor *Fg*Ste3-NLuc protein, the BRET signal was found to increase with galactose concentration, indicating an increase in donor expression and consequently receptor dimer formation, with saturation at 5.6 % galactose. This value, referred to as BRET_max_, indicates the maximum BRET signal reached at receptor interaction saturation on the surface, where each donor interacts with an acceptor. To negate the possibility of detecting false signals due to donor protein overexpression, a BRET_50_ value was calculated. BRET_50_ is indicative of the probability of receptors forming complexes when they are co-expressed at the same time and in the same cell corresponding to the receptor-YFP/receptor-NLuc ratio giving 50 % of the maximal BRET signal reached upon saturation. It describes the relative affinity of BRET partners for each other. Here a BRET_50_ value of 2 % galactose was obtained, which was subsequently used for all BRET assays. This is in contrast to the Negative (Rho) strain (**Table 1**), where no change in receptor engagement or BRET ratio is observed (**Figure 2A**).

**Figure 2.**
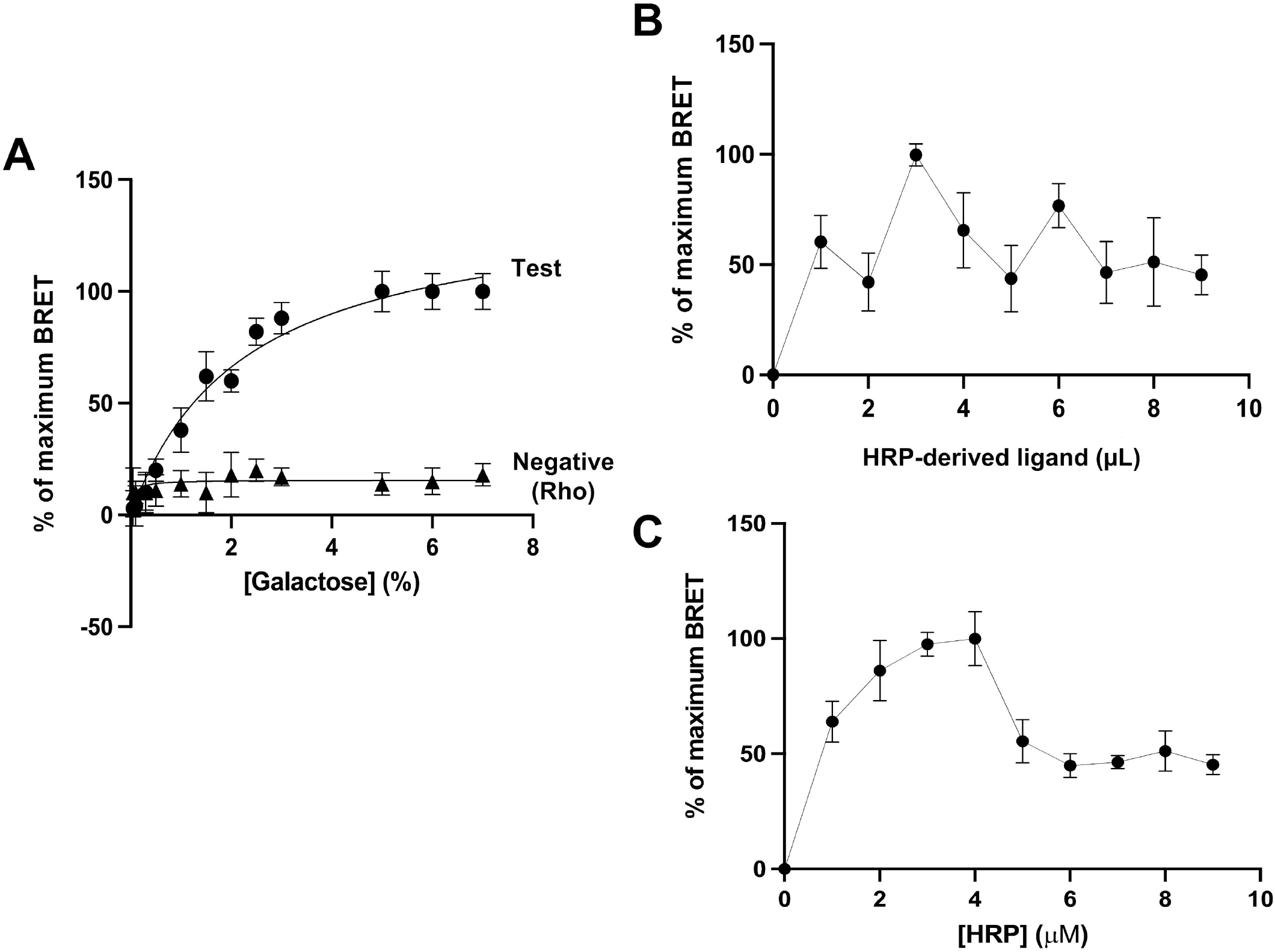
Optimization of BRET assay parameters. **A)** Galactose saturation assay: Galactose saturation assay was carried out at different concentrations of galactose (0-8 %), with the addition of 4 μM HRP, with the Test strain (circles) co-expressing *Fg*Ste3-NLuc and *Fg*Ste2-YFP. The BRET_50_ value was calculated from this to be 2 % galactose concentration, which was used for subsequent experiments. The Negative (Rho) strain (triangles) used here was *Fg*Ste3-NLuc co-expressed with Rho-YFP. **B)** Optimization of HRP-derived ligand application: The BRET response was investigated by treating the Test strain with HRP-derived ligand, by adding increasing volumes of the ligand sample from 1-10μL, with 2 % galactose induction. **C)** Optimization of HRP application: Change in BRET signal was observed over a range of different concentrations of HRP added to the Test strain, with 2 % galactose induction. All data was plotted using Graphpad Prism version 9 (https://www.graphpad.com) and represent the mean and standard deviation of at least two independent experiments done in triplicate.

The optimal concentration of HRP or HRP-derived ligand was also investigated. Different concentrations of HRP-derived ligand or HRP were applied to strain cultures and the effect on receptor-interaction monitored by BRET. In the case of the HRP-derived ligand, while a response was detected indicating the presence of some heterodimer, whether there is any significant concentration-dependence remains unclear (**Figure 2B**). This aspect might be better evaluated once the HRP-derive ligand is characterized and can be better quantified. For HRP, a clear relationship is observed where an increase in HRP concentration leads to an increase in BRET with receptor saturation achieved at 4 μM HRP (**Figure 2C**). Thus, 4 μM HRP was selected and used for all subsequent BRET experiments. Beyond 4 μM, a steep decline was observed that could correspond to dissociation of the heterodimer upon internalization.

### 3.3 Exposure to HRP leads to *Fg*Ste2-*Fg*Ste3 heterodimerization

Based on the optimized parameters obtained above, BRET analysis of the Test yielded a significant BRET signal of greater than 0.1 netBRET (110 mBRET) on exposure to HRP (**Figure 3A and 3C**). This indicates heterodimerization between the *Fg*Ste2 and *Fg*Ste3 receptors, and is in contrast to the low BRET signal from the Negative (Rho) control where the Ste2-YFP construct was replaced with Rho-YFP. The lack of BRET response for the Negative (Rho) strain highlights specificity of the Test strain interaction. An RLU of 5000 for donor and 2000 for acceptor was set as the threshold values above which signal was considered significant, with these conditions being met in the Test, Positive and Negative (Rho) control strains as expected (**Supplemental Figures S1A, S1B and S1E**). The Negative (NR) strain, with no reporter fusions, showed on very low fluorescent signals representing background emissions, as mentioned above, and was subtracted from Test strain and Negative (Rho) strain values (**Supplemental Figure S1C**). To account for background emissions arising from NLuc possibly overlapping into the YFP emission spectra, emissions at 480 nm and 530 nm arising from the *Fg*Ste3-NLuc Donor-alone control strain (**Supplemental Figure S1D**) were used to calculate a Donor-alone BRET ratio which was also subtracted from the values obtained for all other strains.

**Figure 3.**
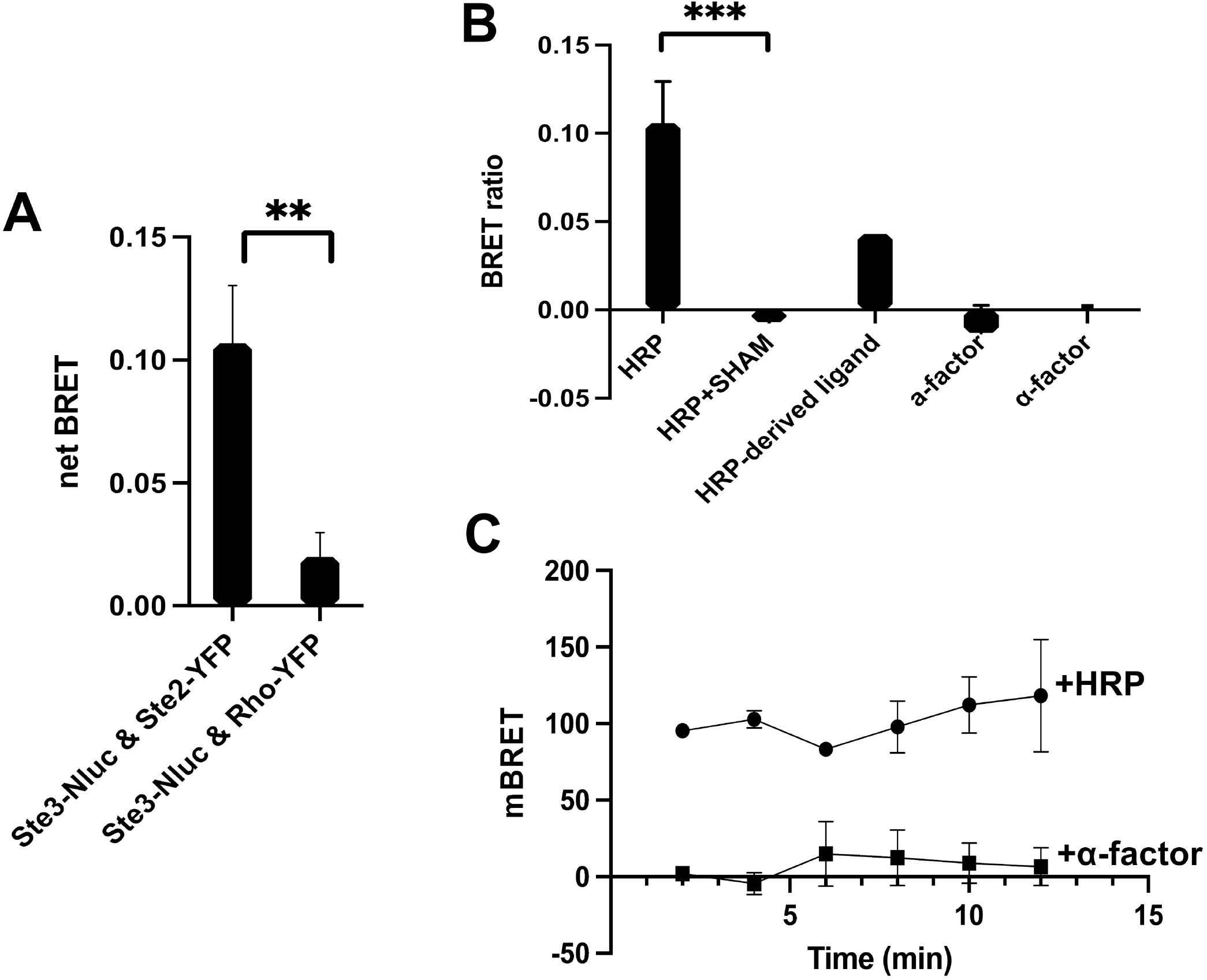
Characterization of *Fg*Ste2-*Fg*Ste3 heterodimer by BRET. **A)** netBRET is calculated for the Test strain: The specificity of the interaction was validated using the Negative (Rho) strain i.e *Fg*Ste3-NLuc was co-expressed with Rho-YFP (**p < 0.01). Data represents the mean and standard deviation of at least two independent experiments done in triplicate. **B)** BRET ratio obtained for the Test strain under different treatment conditions: Final netBRET ratio was calculated for Test strain treated with 4 μM HRP, 4 HRP μM deactivated by SHAM, 3μL HRP-derived ligand, 1 μM Pheromone A and 1 μM Pheromone α (***p < 0.001). Data represents the mean and standard deviation of at least two independent experiments done in triplicate. **C)** Change in BRET with time: milliBRET (mBRET) was calculated for the Test strain, upon induction with either HRP and α-factor pheromone. Each data point represents the mean and standard deviation of ten technical replicates.

Subsequently, the ability of different ligands to stimulate formation of the heterodimer between *Fg*Ste2 and *Fg*Ste3 was investigated (**Figure 3B**). The observed netBRET ratios upon addition of 4 μM HRP were compared to the those arising from treatment with 1 μM pheromones a-factor and α-factor. In fact, no evidence of any BRET response was detected in the presence 0-10 μM pheromones. Furthermore, when HRP was treated with its inhibitor (1 mM SHAM) prior to application to the assay, the observed BRET response was completely lost. Interestingly, 3μL of the HRP-derived ligand gave close to half of the maximum response observed compared to the response upon HRP addition. Together these BRET results emphasize the specificity of the HRP and HRP-derived ligand -induced heterodimerization observed between *Fg*Ste2 and *Fg*Ste3. A lack of any significant time dependence upon application of HRP (**Figure 3C**), suggests that heterodimer formation was already largely saturated by the time the sample was first monitored 5 minutes after adding the substrates. As expected, for α-factor, no significant time dependent BRET response was observed, validating the lack of induction of heterodimerization upon its addition.

### 3.4 Validation of the formation of a peroxidase-induced *Fg*Ste2-*Fg*Ste3 heterodimer by tandem affinity pull-down

To validate *Fg*Ste2-*Fg*Ste3 receptor heterodimerization, a tandem affinity pulldown experiment was performed using the Test, Negative (NR) and Donor-alone strains. After expression and solubilization of the membrane receptors, the obtained detergent containing cell lysates were immunoprecipitated with anti-GFP agarose beads (known to also have affinity for YFP) and then probed for the presence of *Fg*Ste3-NLuc with anti-NLuc antibody by Western blot analysis. One prominent band at approximately 75 kDa representing the monomeric form of recombinant *Fg*Ste3-NLuc was observed for the Test strain (**Figure 4**), compared to the predicted molecular weight of 73,460 kDa. The 75 kDa band was not observed in the Negative (NR) or Donor-alone strains, emphasizing specificity of the antibody and that NLuc does not have any affinity for the Anti-GFP agarose beads itself.

**Figure 4.**
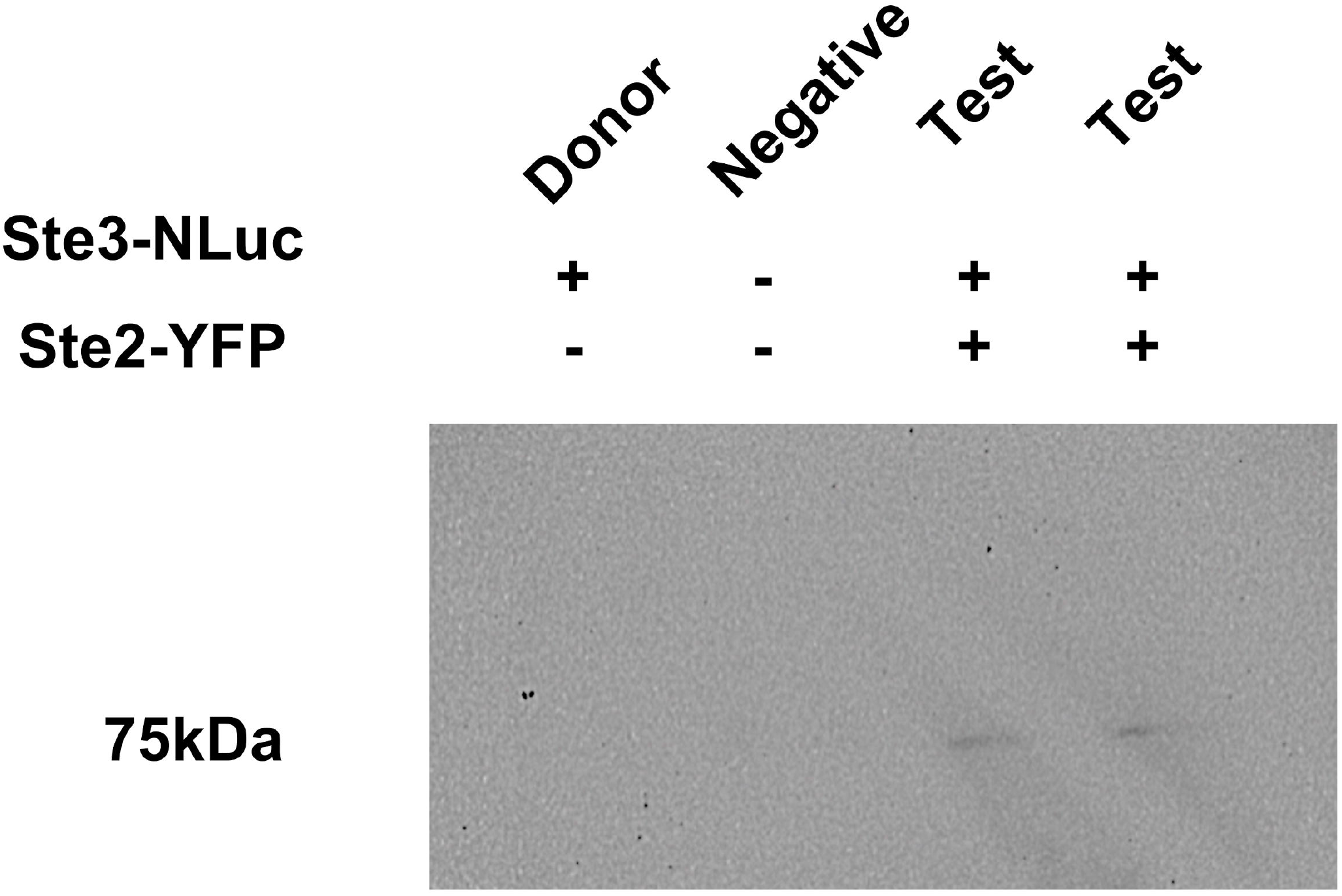
Validation of the *Fg*Ste3 and *Fg*Ste2 interaction by tandem affinity pulldown. The interaction between the receptors in the Test strain was confirmed using *Fg*Ste2-YFP as the bait protein and *Fg*Ste3-NLuc as prey protein. Anti-GFP agarose beads with specificity for YFP were used to pulldown interacting proteins. Western blot was performed using Anti-NLuc antibody.

## 4 Discussion

GPCRs are important signaling molecules that play a crucial role in numerous physiological processes in cells involving responses to external stimuli. GPCRs are known to be activated by the binding of a ligand and are thought to either work independently or form homodimers or heterodimers to initiate downstream signaling events. GPCR heterodimerization has been shown to control a wide range of cellular and physiological processes that differ from the functions of the monomeric versions of the receptors (Calebiro & Godbole, 2018; Sleno & Hébert, 2018). In the plant pathogen *F. graminearum*, we have demonstrated that two GPCRs, *Fg*Ste2 and *Fg*Ste3, are involved in mediating chemotropic responses to host derived peroxidases (Sharma et al., 2022a; Sridhar et al., 2020). Individual deletions of each receptor abolished chemotropic response towards peroxidase, highlighting that each receptor was only active in the presence of the other. Now, through BRET assays and affinity pulldown using a recombinant *S. cerevisiae* system, we have demonstrated the formation of HRP and HRP-derived ligand -stimulated heterodimers between *Fg*Ste2 and *Fg*Ste3. Mechanistically, this heterodimerization could account for the fact that neither receptor can actively mediate chemotropism in the absence of the other. At the same time, at least two different dimer-stimulation models must be considered: ‘ROS-burst mediated’ versus ‘Ligand-binding mediated’.

From the perspective that peroxidase, including HRP, is a notable component of a much larger Reactive Oxygen Species (ROS) system, possible broader roles of ROS are first considered. Common examples of ROS include superoxide anion (·O_2_ −), hydrogen peroxide (H_2_O_2_), hydroxyl radical (·OH) and nitric oxide (·NO). While high concentrations of ROS can cause damage to proteins, lipids, and nucleic acids, low levels of ROS are known to play a critical role in cellular signaling, including the regulation of ion channels, protein phosphorylation, and transcription factors. Thus, the idea of ROS-burst leading to heterodimerization of *Fg*Ste3 and *Fg*Ste2 receptors must be considered (**Figure 5A**). ROS stress is known to modulate membrane microdomains generating spatial proximity of membrane proteins. Major components of the membrane include cholesterols, unsaturated fatty acids, phosphatidyl choline and lipids, all of which are known to be oxidized through action of peroxidases (Eze, 1992; Su et al., 2019). Whether some form of membrane oxidation effect might be contributing to formation or stabilization of the *Fg*Ste3-*Fg*Ste2 heterodimer through modification of lateral membrane pressure on the receptors remains to be determined. At the same time, a large number of studies have shown that many different proteins form dimers in response to oxidative stress (Schieber & Chandel, 2014; Thannickal & Fanburg, 2000). In this context, ROS have been shown to promote formation of disulphide bonds through modification of proteins by oxidation of cysteine residues leading to intramolecular disulphide linkages in the thiol groups (Baba & Bhatnagar, 2018; G. Lee et al., 2016; Miki & Funato, 2012). Notably with relevance to MAPK pathways, Gotoh and Cooper demonstrated ROS dependent homodimerization of Apoptosis signal-regulating kinase 1 (ASK-1) (Gotoh & Cooper, 1998; Hancock et al., 2006; Thannickal & Fanburg, 2000), although whether the mechanism underlying this relates to disulfide formation remains enigmatic. Both *Fg*Ste2 and *Fg*Ste3 have 6 cysteine residues each, located in TM segments and extracellular or intracellular loops supporting this possibility (**Supplemental Figure S2**). However, ultimately the dearth of documented evidence for ROS-burst stimulated GPCR dimerization to date highlights a lack of president for this model.

**Figure 5.**
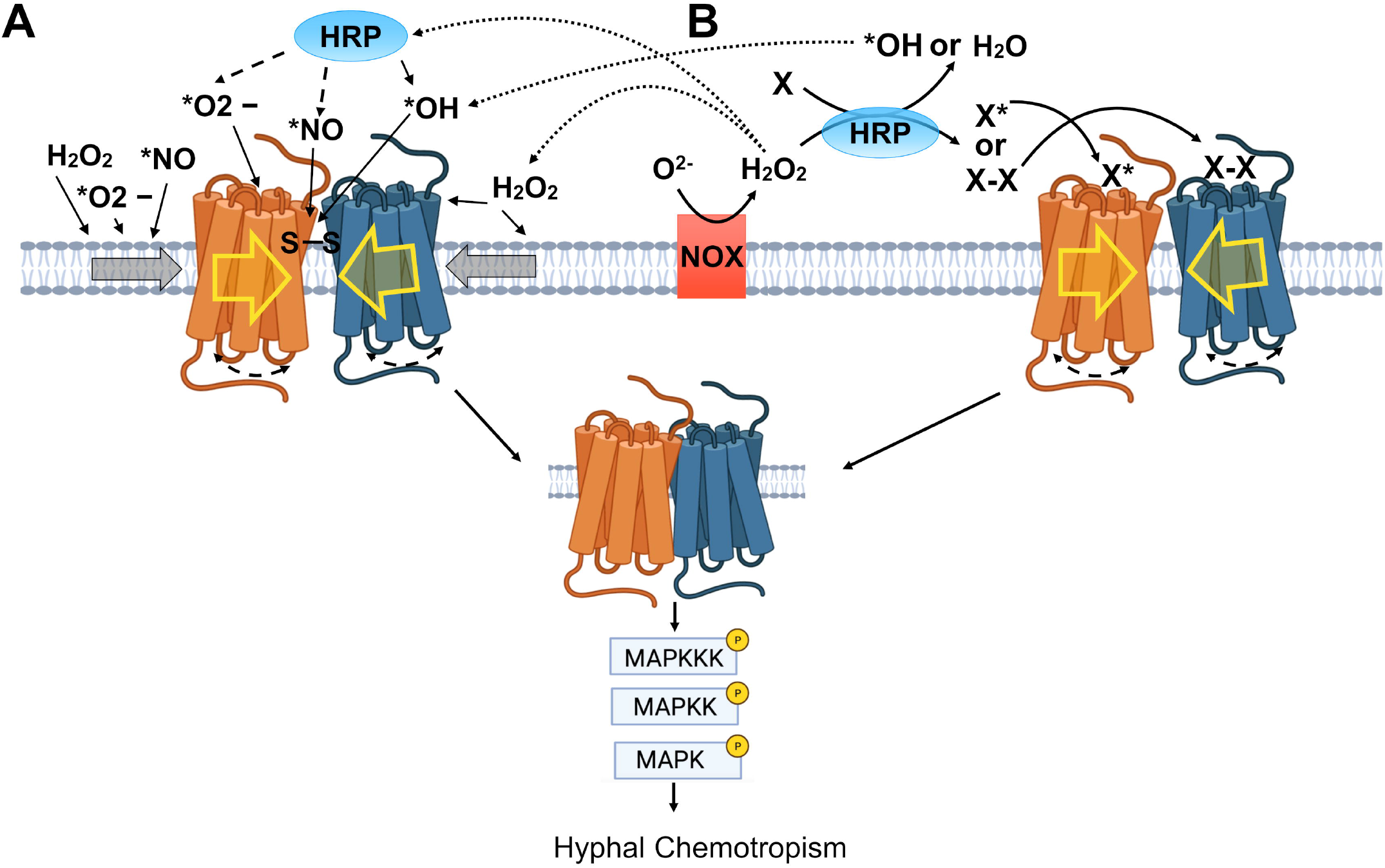
Hypothetical models of heterodimerization of *Fg*Ste3 and *Fg*Ste2 receptors. **A**. ROS-burst mediated model. The formation of this receptor dimer is mediated by the action of peroxidase induced ROS-burst molecules acting directly to oxidize the membrane or surrounding proteins, modifying the environment, changing lateral membrane pressure (grey box arrows) such that the GPCRs are encouraged to form a dimer that may be stabilized by a disulphide bridge. **B**. Ligand-binding mediated model. The formation of this heterodimer relies on the binding of the HRP-derived *Fg*Ste2 and *Fg*Ste3 ligand compound(s), where ligand-induced conformational changes lead to dimerization of the receptors. HRP is a peroxidase that uses H_2_O_2_ as an electron acceptor in the catalysis of a number of oxidative reactions that involve a wide variety of organic and inorganic substrates from fungal cell walls or cell membranes (referred to as X in the figure), one or more of which is anticipated to form the HRP-derived ligand. Generally, peroxidases act on monolignols and sugars. Radicals formed can also induce cross linking (X-X) between cell wall components. The possibility that it is a combination of both models contributing to heterodimerization is emphasized by the arrows going from panel B to panel A.

Thus, we consider an alternate possibility, the ‘Ligand-binding mediated’ model for heterodimerization (**Figure 5B**). Ligand induced heterodimerization has been demonstrated for a variety of GPCRs. For instance, CXCR4 and the δ-opioid receptor form dimer when induced with their respective agonists SDF1α and [d-Pen2,d-Pen5]enkephalin (Pello et al., 2008).

As the ‘ligand’ scenario may pertain to the *Fg*Ste2-<u>Fg</u>Ste3 heterodimerization, ROS, while still involved, is merely a co-factor (H_2_O_2_) enabling production of a ligand via the action of peroxidase (HRP) on fungal cell wall components. Peroxidases can conduct ‘peroxidative’ reactions that polymerize monomeric compounds such as monolignols, into polymeric structures such as lignin. Alternatively, peroxidases can conduct hydroxylic reactions, for example against existing polysaccharide chains, releasing shorter sugar chains or even monomeric sugar units (Passardi et al., 2004). In both cases H_2_O_2_ is consumed as a source of electrons. In this model, the HRP-derived ligands (speculatively, polymerized lignols or small carbohydrate units) are responsible for inducing receptor heterodimerization upon binding to one or both of *Fg*Ste2 and *Fg*Ste3 (**Figure 5B**). Beyond this, ligand-mediated activation and dimerization of the receptors may subsequently lead to up regulation of NOX expression (see below), making more H_2_O_2_ and allowing HRP to make even more ligand, in a feedforward loop.

With respect to the relationship between GPCRs and NOX, in *F. oxysporum*, NOX expression has been shown to regulate chemotropic bending (Nordzieke et al., 2019; Sharma et al., 2022a). Specifically, deletion of NOX genes in *F. oxysporum* eliminated chemotropism, where addition of exogeneous H_2_O_2_ to Δ*noxB* and Δ*noxR* knockouts rescued chemotropic growth. These findings emphasize the importance of these enzymes and H_2_O_2_ in mediating chemotropism. However, H_2_O_2_ did not rescue chemotropism in a Δ*Foste2* strain, suggesting that H_2_O_2_ does not directly activate CWI-MAPK in this system (Nordzieke et al., 2019). Thus, this outcome suggests that the action of H_2_O_2_ is mediated via the receptor, consistent with the second ‘Ligand-binding mediated’ model (**Figure 5B**). At the same time, transcriptomic analysis in WT *F. graminearum* on exposure to HRP shows upregulation of the NOX family gene, 6-hydroxy-d-nicotine oxidase (FGSG_03616), an effect not observed in the *Fgste3*Δ strain. This emphasizes a connection between NOX expression, peroxidase activity and *Fg*Ste3 signalling. Finally, it is interesting to note that beyond *Fusarium*, additional interactions between NOX and GPCRS have been detected. For example, induction of both angiotensin (ANG) II and serotonin (5-HT) by ligand binding was found to induce the activation of a NOX-like enzyme, leading to the generation of intracellular ROS. Accumulation of the ROS, in turn, led to activation of the p38 mitogen-activated protein kinase (MAPK) pathway mediating ANG II dependent cell hypertrophy (Griendling et al., 1994; S. L. Lee et al., 1998). Whether this example of ROS production-MAPK pathway crosstalk is relevant to the case of *Fg*Ste2-*Fg*Ste3 heterodimerization remains enigmatic.

Our experiments testing HRP and HRP-derived ligand indicate a robust BRET response on HRP addition, with a weaker response observed for the ligand. The partial response to the ligand is indicative of an HRP-derived ligand stimulated heterodimer. That no clear concentration-dependence could be established may pertain to our current inability to quantify the HRP-derived ligand, due to the identity of the ligand being unknown at this time. In terms of investigations towards identifying the nature of the ligand, previous investigations from our lab have shown that the ligand being formed is of fungal origin (Sharma et al., 2022a; Sridhar et al., 2020), with the enzyme substrate presented either on the cell wall or cell membrane or both in *F. graminearum*, which are rich sources of peroxidase substrates (Lüthje & Martinez-Cortes, 2018; Schneider et al., 2023). However that HRP-alone can stimulate the *Fg*Ste2-*Fg*Ste3 heterodimer formation in *S*.*cerevisiae* cannot be ignored. This observation would have to be supported either by i) *S. cerevisiae* presenting peroxidase substrates similar to those of *Fusarium*, for the production of ligand, or ii) a more general ROS-burst arising from the peroxidase activity is actually in play. From this perspective, it is interesting to note that low levels of chemotropic stimulating HRP-derived ligand were recently isolated from *S. cerevisiae* (Sridhar, 2023).

Finally, it is interesting to note that other previous reports for these fungal pheromone receptors have highlighted interdependence on (if not direct physical interactions between) each other. In *S. cerevisiae* to enable mating, the diffusible α-(WHWLQLKPGQPMY) and a-(YIIKGVEWDPA) mating factor pheromones must bind their respective *Sc*Ste2p and *Sc*Ste3p receptors at the same time to elicit the mating response (Leberer et al., 1997; Martin et al., 2011; Shi et al., 2007). Moreover, membrane juxtaposition during the later stages of mating has been proposed to be mediated by pheromone-stimulated cross-membrane interactions between Ste2 and Ste3 receptors in *S. cerevisiae* (Shi et al., 2007). In another example, in *F. oxysporum*, the *Fo*Ste2 and *Fo*Ste3 receptors are known to work in an autocrine fashion to control conidial germination in a density dependent manner, where the presence of a-factor can quench α-factor and lead to conidia repression (Turrá et al., 2015; Vitale et al., 2019). However, the ‘classical’ heterodimer formation observed between *Fg*Ste3 and *Fg*Ste2 in response to peroxidase and HRP-derived ligand in this study, is the first heterodimer report for these receptors.

In conclusion, heterodimers form unique signaling complexes augmenting the versatility of physiological responses and signaling pathways activated by GPCRs. At the same time this added level of diversification increases the potential to design novel and more selective and specific drugs and anti-fungal compounds against GPCRs. The *Fg*Ste2-*Fg*Ste3 heterodimer reported here is a new fascinating example of this dimerization phenomenon. The arising hypotheses pointing to the possibility that both the action of peroxidase in the context of ROS-burst activity, as well as the HRP-derived ligand might be needed to obtain a heterodimer that activates the CWI-MAPK pathway cannot be ignored. Further investigations (biophysical and structural) are required to characterize the molecular basis of this heterodimerization, toward novel anti-fungal development.

## 5 Author Contributions

T. S. planned and carried out most of the experiments, analyzed and interpreted the data, and wrote the first draft of the paper. R.J. contributed to designing the BRET experiments and methodology and helped with preliminary experiments. D. Z. and A. M. contributed to confocal microscopy imaging. M.C.L. conceived the idea, obtained the funding, oversaw the planning of experiments, interpreted the data, and helped write the final version of the manuscript.

## 7 Acknowledgements

We are grateful to Drs. John Allingham and Pooja Sridhar (Queen’s University) and Dr. Umar Iqbal (National Research Council of Canada) for discussions. This report represents National Research Council of Canada Communication # 58423.

## 8 Copyright

Copyright (c) 2023 His Majesty the King in Right of Canada.

## 10 Supplementary Material

BRET Supplementary Figures_230822.pdf: includes Supplemental Figures S1 and S2.

## 11 Data Availability Statement

Any data sets and materials are available upon request to interested researchers.

