## Supplemental Figures S1 and S2 for "*Fusarium graminearum* Ste2 and Ste3 Receptors Undergo Peroxidase Induced Heterodimerization"

### Supplementary Figures S1 and S2

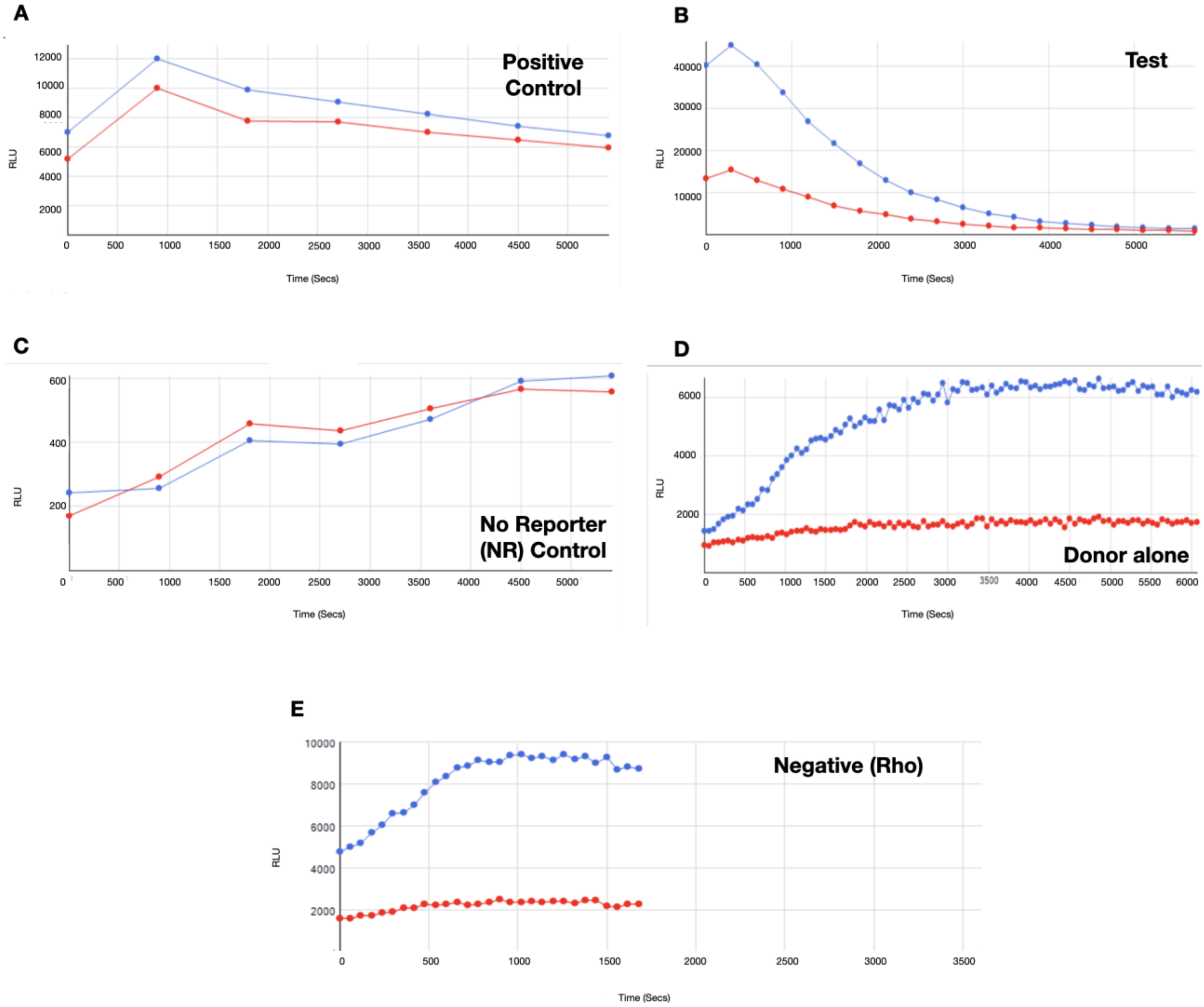


**F**

| | 1 | 2 | 3 | 4 | 5 | 6 | 7 | 8 | 9 | 10 | 11 | 12 |
|---|---|---|---|---|---|---|---|---|---|---|---|---|
| A | 2.0e4 | 2.2e4 | 2.2e4 | 2.1e4 | 2.0e4 | 2.1e4 | 2.5e4 | 2.3e4 | 2.4e4 | | | |
| B | 2.2e4 | 2.0e4 | 2.0e4 | 2.0e4 | 2.2e4 | 2.3e4 | 2.3e4 | 2.5e4 | 2.4e4 | | | |
| C | | | | | | | | | | | | |
| D | | | | | | | | | | | | |
| E | | | | | | | | | | | | |
| F | | | | | | | | | | | | |
| G | | | | | | | | | | | | |
| H | | | | | | | | | | | | |

**Negative (Rho) YFP fluorescence**

**G**

| | 1 | 2 | 3 | 4 | 5 | 6 | 7 | 8 | 9 | 10 | 11 | 12 |
|---|---|---|---|---|---|---|---|---|---|---|---|---|
| A | | | | | | | | | | | | |
| B | | | | | | 1612.4 | 1567.7 | 1588.0 | 1646.3 | 1624.4 | 1712.0 | 1642.5 |
| C | | | | | | 1438.0 | 1394.4 | 1382.9 | 1397.2 | 1402.2 | 1346.9 | 1462.6 |
| D | | | | | | | | | | | | |
| E | | | | | | | | | | | | |
| F | | | | | | | | | | | | |
| G | | | | | | | | | | | | |
| H | | | | | | | | | | | | |

**Negative (NR) YFP fluorescence**

**Supplementary Figure S1: Raw data for each strain used in the BRET study**. Strain samples were treated with 4µM of HRP and then signals recorded. The line in blue represents the Relative Luminescence Units (RLU) for NLuc and red represents the RLU for YFP. BRET ratio was calculated for each of the above strains (A-E). To verify controls, fluorescence was calculated for F) Negative control (Rho) and G) No reporter (NR) negative control respectively, in the 96 well plate.

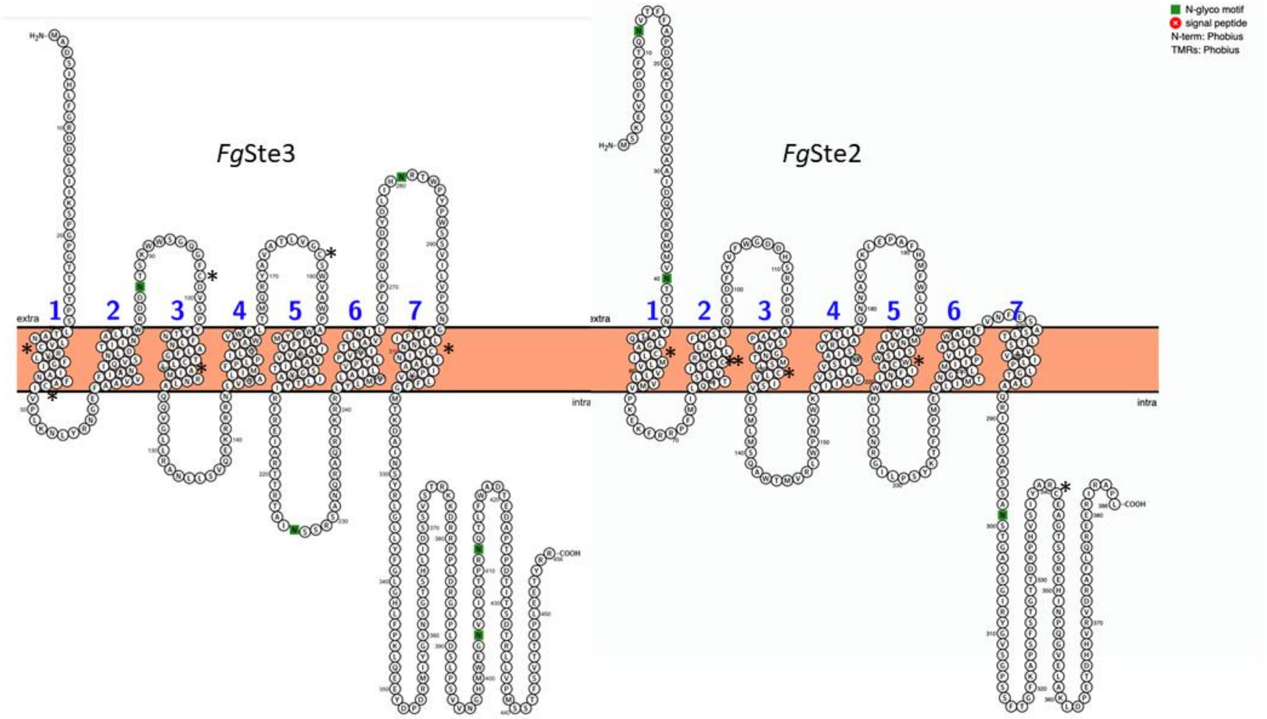


**Supplementary Figure S2. Position of Cysteine Residues in *Fg*Ste2 and *Fg*Ste3 receptors.** Cysteine residues are highlighted by an asterix symbol.
